# ADAPTATION OF DOLPHINS (*TURSIOPS TRUNCATUS*) LOCATION SIGNALS WHEN SEARCHING AND RECOGNIZING OBJECTS HIDDEN BY SEA SEDIMENTS

**DOI:** 10.1101/2021.03.30.437749

**Authors:** K. A. Zaitseva, V. I. Korolev, A. V. Akhi, A. A. Akhi

## Abstract

An experimental study was conducted to research dolphin’s sonar adaptation capabilities for location of objects obscured by marine sediments. It was shown that dolphins are able to alter spectral and temporal characteristics of their location signals. Adaptive change of length, spectral width, and amplitude of impulses provides optimal tools to fight interference and allows dolphins to effectively identify objects.

**SUMMARY:** This work describes dolphin’s search, location and recognition objects hidden by sea sediments and changes in its location signal structure.

## INTRODUCTION

Echolocation is an active process of extracting information about the environment and is one of the perfect tools of nature. The process of evolution led to formation and perfection of this highly adaptive function in many marine and land animals. The main condition for formation of the echolocation is significant limitation on animal’s ability to use vision. Under this condition sound often becomes the primary source of information about the environment. Hearing becomes the lead sensory system. Marine mammals, in particular dolphins, exhibit great development of echolocation.

Multiple experimental studies show high capabilities of dolphin’s echolocator and high efficiency of its locational analysis. Dolphins are capable of distinguishing objects with only 2% difference in size; objects with different shape and material; able to distinguish relative position of objects in a complex target; to locate underwater objects several times smaller than the wavelength of the signal used.

There are a lot of experimental studies that research dolphin’s sonar capabilities. In most of them the location object is placed in the water column. The problem of effective location of underwater objects on the seabed especially the ones obscured by marine sediments is not studied well. This is one of the most relevant problems in modern hydroacoustics. Its solution is tightly connected to the research of the World Ocean shelf, location of small objects like bottom mines and torpedoes, so called aircraft “black boxes”, terrorist devices for attacks on ports and near-sea energy infrastructure objects, conduction of oil and gas works, discovery of defected underwater tubes and cables etc.

Trained dolphins are known to have been used for delivery of cargo, instruments, and information from underwater vehicles or scuba divers to the surface and back. Dolphin’s echolocation ability allows to use them for underwater object search especially for objects at the bottom or in the layer of sea sediments. First experimental studies of dolphin’s ability to locate objects in sediments were conducted after it was noted that dolphins are capable of finding fish burrowed into the sand. In (Rossbach and Herzing, 1997) it is described in high detail how dolphins search for fish hidden in coral sand, dig it up and capture it with rostrum. In (Roitblat et al., 1995) authors developed and implemented bionic location system that used dolphin-like signals. This echolocation system used neural networks and demonstrated ability to locate objects submerged into the bottom silt for up to 10cm. The study shows potential of further research of dolphin’s ability to locate objects at the sea bottom. An experimental comparison between man-made echolocation system and dolphin’s ability to locate object was carried out in (Nachtigal et al., 2000). The objects were submerged up to 45cm into the silt. The animals demonstrated better results than the man-made system. Similar results were obtained by studies conducted on US Navy bases (Moore, 1997; Martin et al., 2005). The results show that trained dolphins are able to detect underwater mines burrowed into the silt more effectively than technical sonars. Dolphin’s ability to locate targets acoustically imitating fish burrowed into quartz sand was studied in (Dahl, 2007). The dolphin was trained to locate targets and bring them to the trainer. The experiments showed that dolphins use step-by-step scan of the sea bottom to locate the targets. The studies mentioned above are phenomenological. They do not focus on what kind of signals a dolphin uses for location, they do not cover how dolphin’s sonar functions when locating objects at the bottom or obscured by the sediments.

The goal of this study is to research location signals emitted by dolphin’s sonar in the process of location and search of underwater objects obscured by sea sediments.

## MATERIALS AND METHODS

The experiments were conducted on two adult dolphins which have been used in previous acoustic studies. The experiments took place in a pile-net aviary of a sea bay 10×10×6m in size. The bottom soil consisted of a mix of sand, seashells, and silt. The technique of motor food conditioned reflexes was applied during free swimming of the animals. The experiment procedure consisted of two phases. During the first one, the animals were taught to distinguish positive and negative targets. The positive target was a hollow brass cylinder 120mm in height, 100mm in diameter with 5mm thick walls. The negative target was a steel cylinder of the exact same size. Both targets were coloured same to avoid any visual distinction. The targets were presented to animals consequently. For finding the positive target dolphins got a fish. Reaction on the negative target was not reinforced.

Target position was randomly changed. To study dolphin’s ability to locate targets in sediments the study had to be adjusted. To prevent dolphins from using visual or acoustic signals during the target setup the dolphins were taught to swim into a special wooden structure. When given a signal the dolphin would swim into the structure inside the aviary. The structure had a special rostrum-shaped notch. The dolphin placed its rostrum into the notch and awaited motionless for the next signal. The dolphin was placed to be facing 180 degrees away from the target. After the target setup the dolphin was given a signal to leave the structure, assume starting position in the aviary, and start the search. Correct identification of the positive target was indicated by the dolphin touching it with its rostrum and a following hit of an indifferent foam manipulator. After correct identification of the positive target the dolphin got a fish. The experiments were conducted with targets being submerged at the depth of 1,5 meters, placed at the bottom, half-submerged, and submerged 10cm into the sediments. To avoid conditional reaction to a location, the aviary was split into square sectors and each time a sector was chosen randomly for the target placement. After each experiment, the dolphin went back to the starting position.

A receiving hydrophone was places 0,5m away from the target. It was submerged 1cm into the sediments. Echolocation signals were recorded from the start of the experiment all the way till the end. The receiving hydrophone provided a non-uniformity of the amplitude-frequency characteristic of no more than 3 dB in the 0.5-200 kHz band. The signals were fed through an amplifier to an ADC and a computer for recording and subsequent processing. Statistical analysis was carried out using the data processing package (SPSS v25 for Windows). 1500 probing impulses were processed (program “Power graph”).

## RESULTS

Fig. 1 shows characteristic temporal realizations of the probing signals, their energy spectra and accumulation in 3D sweeps of the energy spectra in time at the beginning of the search. The dolphin starts the search of the target in the water column. It uses short (10-12μs) pulses with broadband energy spectrum (up to 140-170 kHz). During the search of the target placed at the bottom, the spectral-temporal parameters of the location signals change (Fig. 2b, 2c). In the bottom area echolocation is accompanied by bottom reverberation. To combat it the dolphin shifts the energy spectrum of the probing pulse towards lower frequencies down below 100 kHz with the maximum around 50 kHz. The pulse duration increases to 14-15μs. Further shift in the probing pulse structure is observed when locating objects are submerged into the sediments (Fig. 2d, 2e). The effective band of the energy spectrum narrows down below 80 kHz, its maximum shifts to the low-frequency region of 20 kHz and increases in amplitude by one and a half times in comparison to the signals recorded during the search at the bottom and four times in comparison to the signals emitted during the search in the water column. The pulse duration also increases to 15-20μs. The pulse duration and the spectral bandwidth were measured at a threshold of 10% of the signal amplitude relative to the noise level. Statistical processing of the obtained experimental data showed the following result:

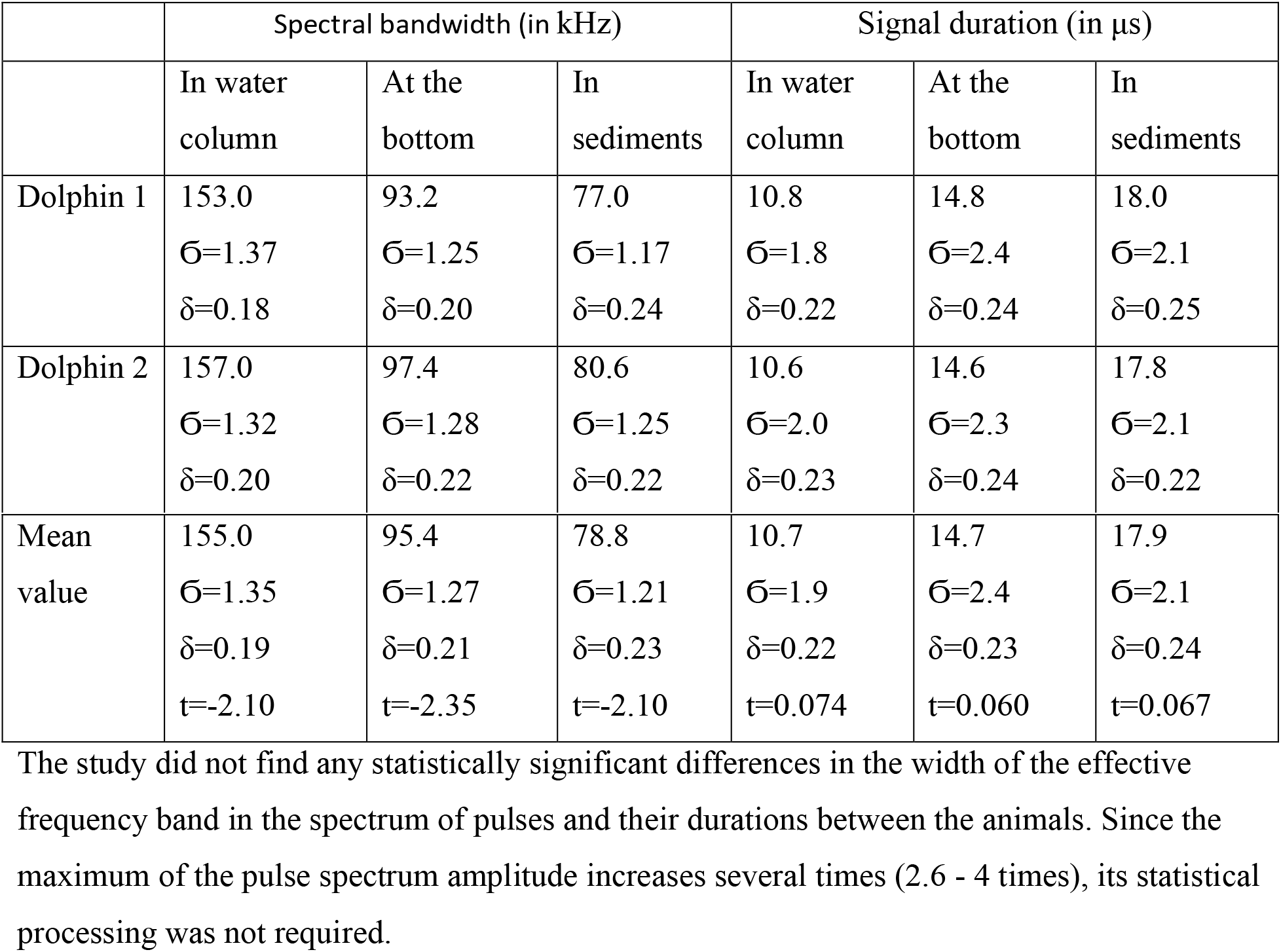

**Fig 1.**
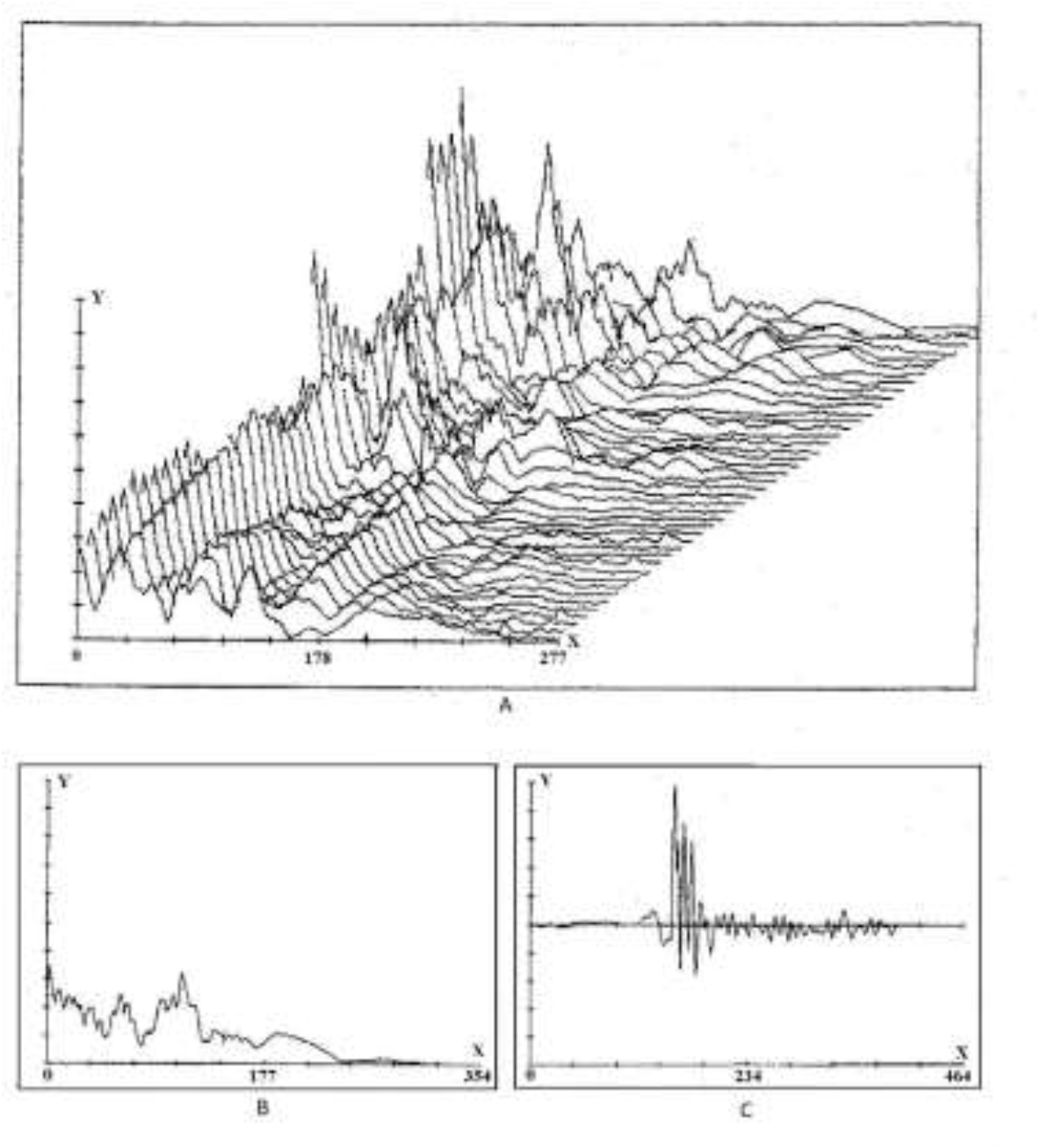
Location signals during search of a target located in the water column. A. accumulation of spectral sweeps in 3D, abscissa – signal frequency in kHz; B. signal spectrum, abscissa – signal frequency in kHz; C. temporal realization of the signal, abscissa – time in μs. The ordinate on A, B, C is the signal amplitude.

**Fig 2.**
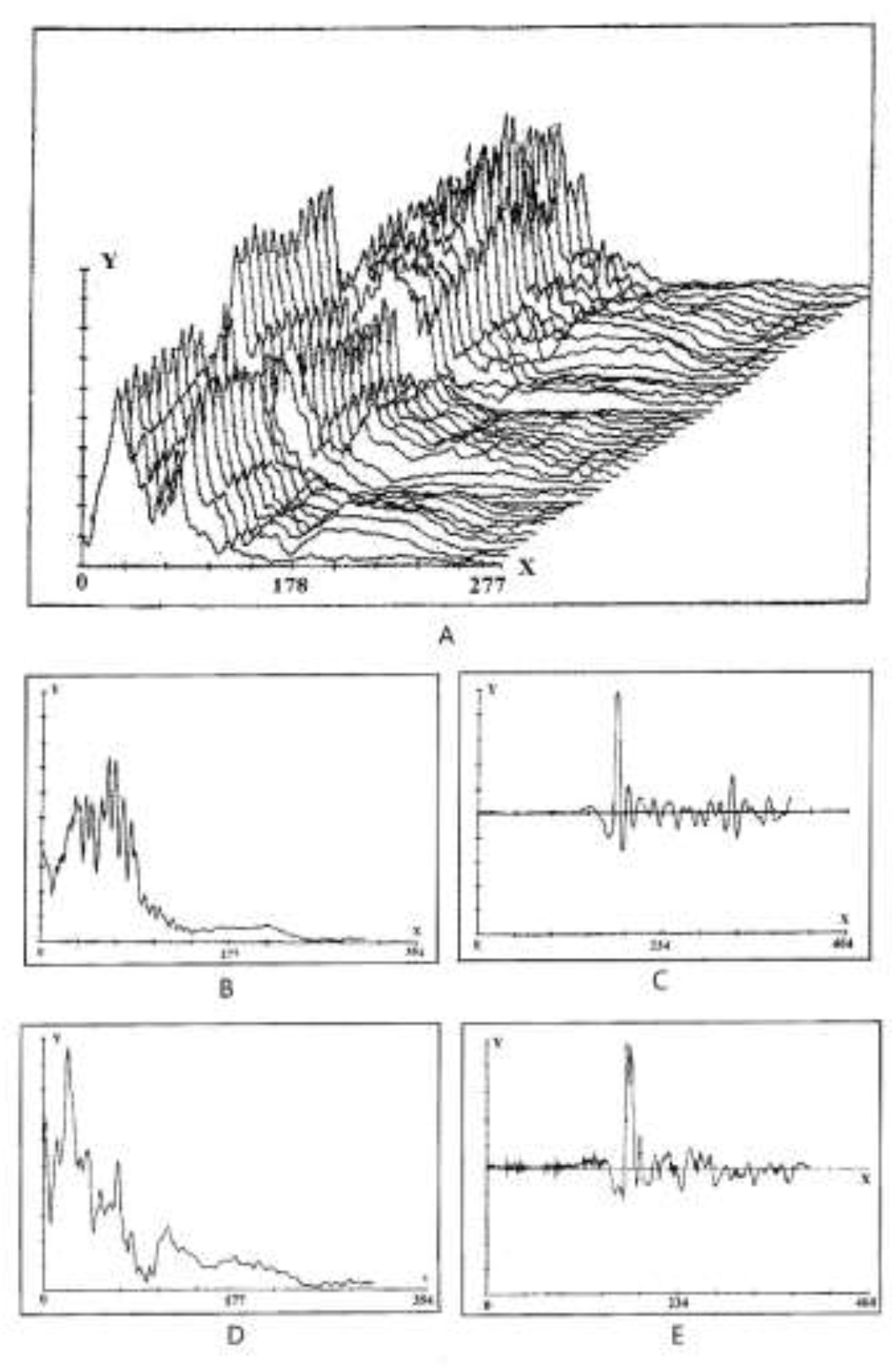
Location signals for near-bottom search, detection, and identification of a target buried in the sediments. A. accumulation of spectral sweeps in 3D from the beginning of the bottom search to identification of the target, abscissa - the signal frequency in kHz; B. signal spectrum during search and detection of a target at the bottom, abscissa - frequency in kHz; C. temporal realization of the signal during search and detection of a target at the bottom, abscissa - time in μs; D. signal spectrum during detection and identification of a target in the sediments, abscissa - frequency in kHz; E. temporal realization of the signal during detection and identification of a target in the sediments, abscissa - time in μs. The ordinate on A, B, C, D, E is the signal amplitude.

## DISCUSSION

The conducted experimental studies showed that dolphin’s locational search is a complex and complicated process. During this process spectral temporal parameters of probing signals change adaptively from the start of the search all the way to the target identification.

Our previous work (Zaitseva et al., 2018) experimentally showed that the variability of the spatial location of the target in the water area has little effect on the efficiency of its detection by the dolphin. The animal locates the target with high probability (78-95%) regardless of target’s location - close to the water surface, in the water column, at the bottom or in the sediments. This fact leads to assumption that a dolphin is capable of adaptively changing the way its acoustic system works and, possibly, changing parameters of the probing signals to achieve reliable reception of the reflected signal. In (Zaitseva et al., 2019), we experimentally investigated how the spectral-temporal structure of the probing pulses changes during the transition from a location-based search for a target in the water column to a search in the near-surface layer. A decrease in the upper frequency, an increase in the duration and the amplitude of the pulses were recorded.

It is known that subtle mechanisms of adaptive changes in the work of dolphin’s sonar system are revealed when animals solve complex acoustic problems. Search, detection, and identification of objects obscured by marine sediments are a challenge for dolphin’s echolocation system. The main difficulties arising in this case are presence of strong bottom reverberation, volumetric reverberation in the ground, sound reflection by the uneven water-ground surface when on the way to the target and the ground-water surface after reflection from the target, attenuation and scattering of the signal on acoustic inhomogeneities of the ground, nonlinear interaction of roll waves in the ground. To overcome these difficulties the dolphin is using low frequency probing signals which are less affected by noise and attenuation on ground inhomogeneities. A significant increase in the maximum amplitude of the signal in combination with an increase in its duration leads to an increase of its energy. The high pulse energy, which provides an increase in the signal-to-noise ratio, allows penetration of the sediment material with numerous inhomogeneities.

It is controversial whether dolphins carry out targeted changes in the spectrum of emitted pulses. Experimental studies that were carried out in 70s with the help of broadband receiving and analyzing equipment did not allow authoritative researchers to come to an unambiguous conclusion on whether any adaptive regular changes in the spectrum of the echolocation pulse exist. Some authors concluded that dolphins lacked purposeful adjustment of spectral composition of location signal and that observed changes were most likely a result of incorrect interpretation of the data or insufficient statistics.

However, around the same time a number of researchers gave a positive answer to this question and provided a confirmation with experimental data on the shift of the maximum in the spectra of location signals in both directions from the maximum of the energy spectrum with the offset reaching 20 kHz or more. To date, a fairly large amount of experimental material exists indicating that when solving a number of location problems a dolphin can use such an adaptation method as spectral detuning. Most often it is used in the presence of interference of various origins. The energy maximum in the spectrum of dolphin’s location signal can shift to the region of lower frequencies (35-50 kHz) with the high-frequency interference of 80 kHz. The presence of a 30kHz tone signal or a pulse with a filling frequency of 30 kHz and a sound pressure of 1200Pa in the noise causes a shift in the spectral maximum to the high frequency region of 65-85 kHz. A dolphin shifts signal spectrum to a frequency range at which the interference intensity is much less than the intensity of the location signal, i.e. tunes in frequency. The shift of the spectrum maximum of the probing pulses to the low-frequency region during repeated presentation of the object was observed in (Au at al., 1974; Au and Snyder, 1980).

Adaptive changes in the duration and the intensity of the location signals emitted by a dolphin in solving problems of varying degrees of complexity have been established in many experimental studies. These changes can be quite significant. According to some data the amplitude of the probing pulses can increase several times. The mechanism for blocking interference by increasing the radiated signal level is well known in both aquatic and terrestrial mammals whether they use acoustic signals for communication or location. Humans use voice intensity adjustment during speech communication in various acoustic conditions (ranging from scream in presence of high noise to whisper in silence). Moreover, a human reacts to the introduction of interference not only by increasing the volume of speech but also by stretching the vowels. The possibility of adaptive changes in the duration of dolphin’s location pulses is limited by its structure. Dolphin’s echolocation signal is a short pulse with a steep edge and a wide spectrum. Pulse characteristics of the signal enable sonar’s high resolution and high efficiency during target search and location. However, the duration of the location pulses is small and is limited. Increase in signal energy is possible through significant increase in its intensity. Similar results were achieved in (Zaitseva et al., 2018) when dolphins search for objects close to the surface in the presence of surface reverberation. It was shown that when a target located at the water-air interface is detected, the perception of the reflected location signal becomes difficult, but the probability of correct detection is quite high. At the same time signal spectrum is narrowed down to 40-60 kHz and its amplitude doubles. A duration increase up to 45μs, narrowing the spectrum, an amplitude increase provide detuning from interference including reverberation and effective identification of the object by increasing the signal energy.

The data obtained in this work indicates that dolphin’s sonar can adaptively simultaneously change three parameters of the emitted signal: duration, frequency, and intensity, enabling search, detection, and identification of an object in the sediments. A dolphin can change both location signal parameters and its own trajectory of movement. We showed in (Zaitseva et al., 2020) that a dolphin is able to change the search strategy, moving from linear motion to the target to complex maneuvering. This allows it to find the optimal position and location angle. Finding optimal conditions is necessary to obtain comprehensive information about the object in the ground. When moving towards the target the dolphin makes rotational movements. The end of its rostrum follows an elliptical trajectory. The shape and spectrum of the locating pulse in this case substantially depend on the angle relative to the longitudinal axis of the animal’s body. By varying this angle, the dolphin irradiates the investigated target with different pulses. The larger the angle, the more low-frequency the pulse spectrum is. As the most difficult task, target identification forces the dolphin to perform complex circular funnel-shaped maneuvers over the object. A circular motion along the surface of the cone with the apex at the target point helps to effectively probe the target at different angles of location and choose the optimal directions to overcome the bottom reverberation. Long-term maneuvering allows to increase the decision-making time required for multiple sounding of the object without losing acoustic contact with the target.

The studies have shown the possibility of adaptive changes in the spectral-temporal parameters of the dolphin sonar depending on the location conditions. Unlike live sonar, technical sonar systems are characterized by predetermined unchanged parameters, in particular narrowband, and cannot quickly adapt to changing conditions.

## COMPLIANCE WITH ETHICAL STANDARTS

All applicable international, national and institutional principles of handling and using experimental animals for scientific purposes were observed. The study did not involve human subjects as research objects.

## AUTHOR CONTRIBUTIONS

Zaitseva K. A. – theory, discussion.

Korolev V. I. – methodology, experiments.

Akhi A. V. – literature review, discussion, technical support.

Akhi A. A. – data handling, technical support.

